# An anti-Galectin-3 intrabody improves primary and memory response of antigen-specific CD8^+^ T cells during viral infection

**DOI:** 10.64898/2026.09.16.751996

**Authors:** Yuviana J Singh, Sharvan Sehrawat

**Author notes:** Correspondence to: Dr. Sharvan Sehrawat.

## Abstract

Genetic ablation approaches are routinely used to establish gene function, but these are not beyond limitations. The intracellularly expressed camelid single-domain antibodies (sdAbs), or nanobodies, can specifically disrupt molecular interactions and, therefore, help ascertain their role in the process of cellular differentiation. The approach has not been evaluated in differentiating T cells or the stimulating DCs during viral infection. We modified the intracellular response of Galectin-3 (Gal-3) using an α-Gal-3 intrabody (IB) in antigen-specific CD8^+^ T cells and assessed their differentiation patterns during a resolving Influenza A virus encoding SIINFEKL epitope (IAV: WSN-SIINFEKL). The α-Gal-3 IB expressors, compared to the cells genetically depleted of Gal-3, exhibit a significantly prolonged activation, enhanced proliferation, and elevated cytokine production following *in vitro* stimulation. The α-Gal-3 IB expressing cells show enhanced functionality and generate an efficient effector response, favoring the generation of memory precursor effector cells (MPECs: CD127^hi^KLRG1^lo^) over short-lived effector cells (SLECs: CD127^lo^KLRG1^hi^) in IAV-infected animals. Such a differentiation pattern results in improved differentiation of effector memory (CD44^hi^CD62L^lo^CCR7^lo^) cells amongst the α-Gal-3 IB-expressing cells. The generated memory cells are efficiently recalled upon challenge infection with MHV68-SIINFEKL and help control the infection better. Therefore, modifying the intracellular response of Gal-3 using a sdAb could improve the primary and memory response of antigen-specific CD8^+^ T cells.

**Author Summary:** Genetic depletion of Gal-3 significantly increases the response of antigen-specific CD8^+^ T cells; whether this effect is attributable to exogenous or endogenous Gal-3 remains to be adequately elucidated. Using an intracellularly expressed α-Gal-3 sdAb that modulates the levels of bioavailable Gal-3, we study the differentiation pattern of CD8^+^ T cells during Influenza A virus infection. The α-Gal-3 intrabody (IB)-expressing CD8^+^ T cells, compared to their control counterparts, generate functionally superior effector and memory responses, and the recalled memory cells help control the infection better. The study establishes the utility of an α-Gal-3 IB as a potential strategy to improve antigen-specific primary and memory response of CD8^+^ T cells.

## Introduction

Galectins (Gals) are evolutionarily conserved β-galactoside-binding proteins that mediate vital cellular functions such as recognition of pathogens, regulation of signaling pathways, and modulation of immune responses (1–3). Gal-3, the only chimeric type galectin, is composed of a carbohydrate recognition domain, i.e., CRD, and a non-lectin, long and flexible N-terminal domain (NTD) (4,5). Gal-3 influences several pathways and physiological processes, such as apoptosis, cellular growth, angiogenesis, and mRNA processing (6–8). While the extracellular Gal-3 predominantly mediates cell adhesive functions by forming lattices with ligands, the intracellular Gal-3 affects cellular survival, proliferation, activation and differentiation by altering the gene expression patterns (9). Owing to its high propensity to bind glycans, Gal-3 interacts with different cell surface expressed molecules, such as the components of the TCR complex, CD45, as well as some of the costimulatory or inhibitory molecules, such as CTLA-4, LAG-3, during the immune synapse formation (10). It has been shown that the genetic removal of Gal-3 significantly improves effector functions of antigen-specific CD8^+^ T cells (8). How intracellularly expressed Gal-3 regulates T cell responses remains to be established. By interacting with the Alix protein, Gal-3 might regulate signaling via TCR (11). Gal-3 localizes at the immunological synapse when naive or memory CD8^+^ T cells undergo activation (12). Furthermore, CD8^+^ T cells from Gal-3 KO mice generated a robust antigen-specific response compared with those from wild-type (WT) mice.

Genetic knockdown approaches or small-molecule inhibitors are used to investigate the role of Gal-3 in pathophysiological processes. Such approaches, however, often have several nonspecific and unaccounted-for effects on cellular processes and pathways (13,14). These include off-target effects or the induced compensatory mechanisms that can either mask or overestimate the observed phenotype, especially when the encoded product acts in multiple pathways (15).

Furthermore, the expression kinetics of different proteins can alter the observed phenotypes, which might be missed with the complete removal of the encoding genes (16,17). Therefore, modifying the bioavailable levels of proteins could unravel less-appreciated or altogether unknown functions in cellular physiology (18,19). This is particularly relevant in defining a protein’s function in the cells of the adaptive immune system, such as T cells. The developmental and differentiation processes of T cells are governed by at least two distinct, sequentially operating programs. The initial programming occurs during the selection process in the thymus, while the other program drives their response patterns following antigenic stimulation or bystander activation in the periphery (20). Therefore, a complete ablation of the gene might confound the interpretation of the obtained results. Although small molecules are used to inhibit or promote protein function, they also exhibit nonspecific effects. Therefore, alternative approaches that modulate the bioavailable levels of macromolecules could be valuable; one such approach is the use of antibodies that interact with the ligand with pinpoint precision (21,22). The variable region of camelid ‘heavy chain only antibodies’ (V_H_H), also referred to as nanobodies or single domain antibodies (sdAb), can be efficiently expressed intracellularly as functional products to disrupt specific functions of proteins by blocking their binding sites (18,23).

Here, we modified the response of Gal-3 in antigen-specific CD8^+^ T cells with the intracellularly expressed anti-Gal-3 intrabody (α-Gal-3 IB) and tracked their fate during the primary and memory phase of a resolving IAV infection. We showed that α-Gal-3 IB expressors mounted enhanced effector and memory responses compared with control cells and better controlled a secondary infection. Such differences were qualitatively different from those observed following genetic knockout of the Gal-3 response.

## Results

### α-Gal-3 intrabody modifies Gal-3 response in the stimulated CD8^+^ T cells

The sequence encoding an α-Gal-3 nanobody and a c-Myc tag was cloned upstream to GFP into the pLenti-GFP vector using the Xba-1 and Bam-H1 restriction sites (Fig. S1A-C). In brief, the sequence of the α-Gal-3 nanobody was amplified using specific forward and reverse primers, and the amplicon was cloned into the pLenti-GFP vector (24) (Fig. S1B). The ligated product was transformed into the Stbl-3 strain of *E. coli*. The PCR-positive colonies were confirmed by restriction digestion using *Sal-1* and *Nde-1* enzymes, yielding a product size of 1606 bp (Fig. S1C). HEK293T cells were then transfected with either the α-Gal-3 nanobody construct or the GFP control vector, and the cells were collected 72 hours post-transfection for analysis. The prepared cell lysates were resolved by SDS-PAGE, and the polypeptides were transferred to PVDF membranes for probing with the anti-c-Myc monoclonal antibody. While the vector control transfected cells did not show any reactivity, a band of ∼47 kDa was detected only in the α-Gal-3 IB-expressing cells (Fig. 1A, upper panel). Anti-GAPDH monoclonal antibody probed membranes showed equal loading of the samples (Fig. 1A, lower panel). The expressed α-Gal-3 intrabody (IB) was pulled down using recombinant 6x histidine-tagged Gal-3. Two polypeptide bands of 30kDa and 47kDa corresponding to the molecular mass of Gal-3 and α-Gal-3-IB fusion products, respectively, were evident in the stained gels (Fig. 1B). pLenti-GFP containing α-Gal-3-IB sequence along with four additional plasmids, i.e., pMD2.G, which contains the sequence for VSV-G, pCMVR.74, Tat1b, and Rev1b were used to transfect HEK293T cells to generate pseudovirus particles encoding the α-Gal-3 single domain antibody. The Ova-specific CD8^+^ (OT1) T cells transduced with the pseudoviruses were FACS-sorted based on their GFP expression (Fig. 1C-E). To confirm the expression of α-Gal-3 IB, cell lysates were prepared, and the resolved electroblotted polypeptides were probed using anti-c-Myc monoclonal antibody. The expression of the fusion product of α-Gal-3 IB was revealed by a band of 47 kDa (Fig. 1F).

**Fig 1.**
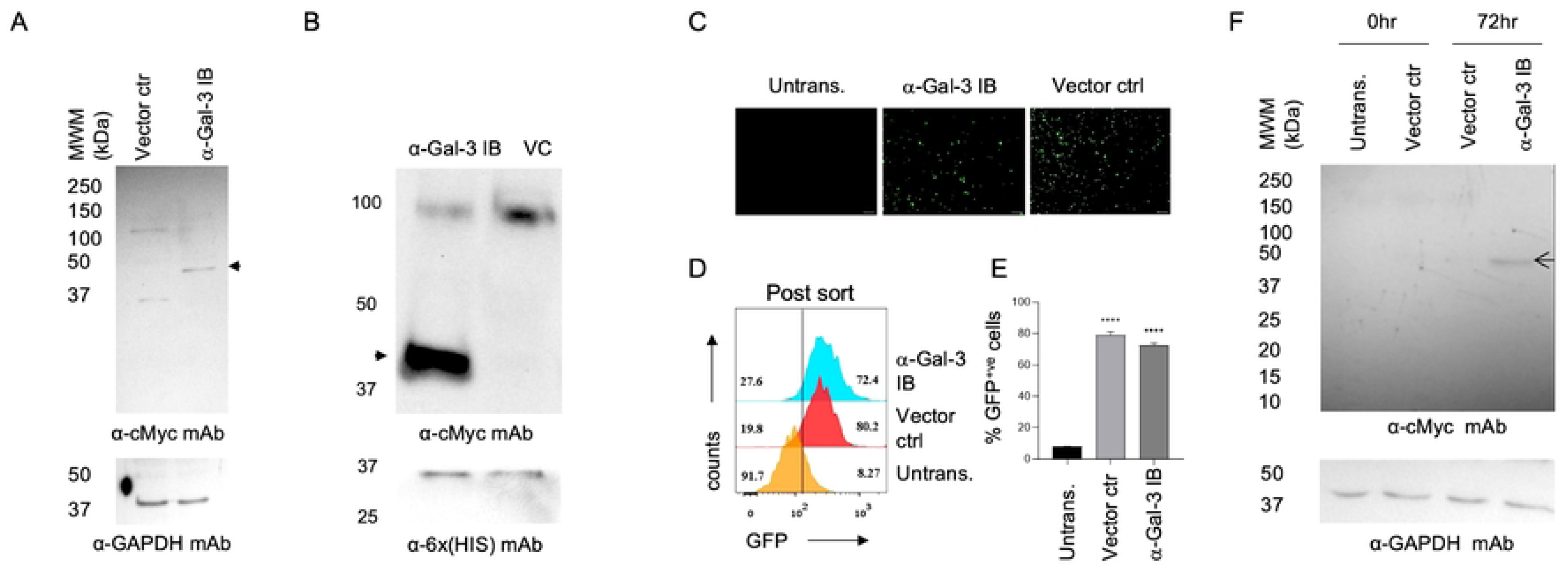
Expression of anti-Gal-3-intrabody (α-Gal-3-IB) by naïve CD8^+^ T cells. A. The HEK 293T cells were transfected with α-Gal-3-IB and the vector control plasmids. The transfected cells were collected after 72 hours and were used to prepare the cell lysates. The cell lysates were run on an SDS-PAGE gel and transferred to a PVDF membrane, which was further probed with an anti-c-Myc monoclonal antibody. This successfully detected the IB by producing a band at approximately 47 kDa. The cell lysates were also probed with anti-GAPDH monoclonal antibody to ensure equal loading of the samples. B. The cell lysates were also used to perform a pull-down experiment to confirm the specificity of the IB for Gal-3. The IB specifically interacted with the immobilized Gal-3, as bands corresponding to Gal-3 and IB (i.e., 30 kDa and 47 kDa) were visible after the membrane was developed with the specific antibodies. C. The α-Gal-3-IB and vector control (VC) expressing pseudoviruses were prepared using the five-plasmid system, and the pseudoviruses were used to transduce naïve CD8^+^ T cells. The transduced CD8^+^ T cells were sorted based on GFP expression. The fluorescent images were captured with an 80 μm scale bar after 72 hours. D. Representative offset histograms from different conditions are shown. The vertical dark solid line separates GFP^+ve^ from GFP^-ve^ populations. E. The bar graphs obtained from multiple experiments are shown as cumulative data. The levels of statistical significance were measured using One-way ANOVA and are shown as (****; p < 0.0001, ***; p < 0.001, **; p < 0.01, *; p ≤ 0.05, and ^ns^; p>0.05. F. The untransduced CD8^+^ T cells, GFP^+ve^ cells from the control vector-transduced, and the anti-Gal-3 IB-expressing condition were sorted to prepare cell lysates. The resolved polypeptides were transferred to PVDF membranes and probed by Western blotting to confirm IB expression in naive CD8^+^ T cells. Anti-c-Myc monoclonal antibody confirmed IB expression in naïve CD8^+^ T cells, and an anti-GAPDH mAb was used to assess equal loading.

Does α-Gal-3 IB modify Gal-3 response in the transduced CD8^+^ T cells? Splenocytes from OT1 TCR transgenic mice, transduced with the pseudoviruses encoding the α-Gal-3 IB, were activated *in vitro* with 0.1 and 1 μg/ml of SIINFEKL peptide (Fig. S2A). After 48 hrs of activation, the α-Gal-3 IB expressing CD8^+^ T cells showed significantly lower frequencies of Gal-3 positive cells in comparison to those transduced with control pseudoviruses (42.13 ± 1.89% vs 54.33 ± 2.98 %) at the stimulating concentration of 1μg/ml (Fig. S2B and C). A similar reduction in Gal-3 levels was observed in culture supernatants, as measured by ELISA (Fig. S2D). Gal-3 occupancy by the expressed α-Gal-3 IB could have rendered intracellular detection inefficient. Furthermore, the antibody could have channeled the immune complexes for degradation, or it interfered with the release of Gal-3 via a non-canonical secretory pathway. Taken together, we show that the α-Gal-3 IB altered Gal-3 response in the stimulated CD8^+^ T cells.

To directly assess the interaction of α-Gal-3 IB and the produced Gal-3 within stimulated CD8^+^ T cells, we performed colocalization experiments. Control and the α-Gal-3 IB expressing OT1 cells were stimulated via their co-receptors using anti-CD3/28 antibodies and analyzed for colocalization of α-Gal-3 IB, as detected by GFP, Gal-3, and Zap70, a kinase recruited to the cytoplasmic domain of the TCR complex to facilitate the cellular activation process (10,25). Both Gal-3 and the α-Gal-3 IB showed significantly higher colocalization with a Pearson’s coefficient (r) value of 0.75 compared to the vector transduced control cells, which showed an (r) value of 0.35 (Fig. 2A and B). While the control cells showed a colocalization coefficient of Gal-3 and Zap70 of 0.73, the value was 0.59 for α-Gal-3 IB-expressing cells, indicating a sequestration of Gal-3 by the expressed anti-Gal-3 IB (Fig. 2A and B). Interestingly, α-Gal-3 IB expressors compared with control cells showed a diffused staining for Gal-3 and a more efficient localization of Zap70 at the periphery of cells, indicating enhanced activation following their coreceptor ligation (Fig. 2A and B). Alix, a known interactor of Gal-3, is involved in TCR signaling (25). While unstimulated cells show poor co-localization of Alix with Gal-3, following stimulation, Gal-3 co-localizes more efficiently with Alix, with a colocalization coefficient of 0.74 (Fig 2C and D). The α-Gal-3 IB-expressing cells showed reduced colocalization of Gal-3 with Alix, with r values of 0.60 (Fig 2C and D). Alix co-localized with the α-Gal-3 IB more efficiently in the stimulated cells, with the r values of more than 0.80 (Fig 2C and D). These results, therefore, suggest that expressed α-Gal-3 IB colocalized with Gal-3 in the stimulated CD8^+^ T cells and, in so doing, induced their efficient activation.

**Fig 2.**
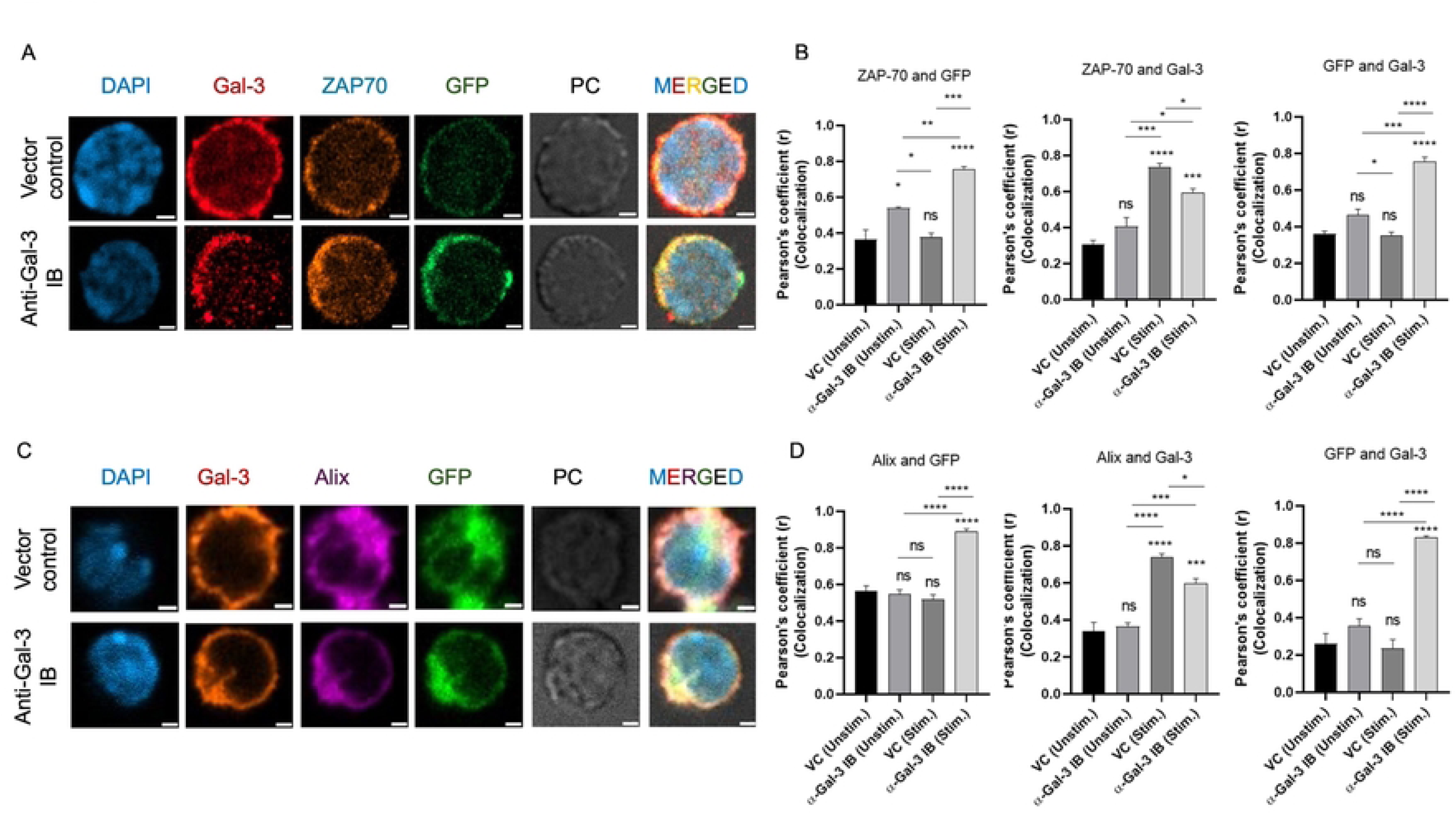
α-Gal-3 IB colocalizes with intracellular Gal-3 and induces efficient activation in the stimulated CD8^+^ T cells. A-D. Spleen and lymph nodes were pooled from C57BL/6 mice in 10% RPMI, and CD8^+^ T cells were MACS-purified. The purified CD8^+^ T cells were then spin-transduced with a pseudovirus expressing an anti-Gal-3 sdAb (IB) or a vector control (VC). After 72 hours, GFP-positive CD8^+^ T cells were analyzed and FACS-sorted for performing confocal microscopy. The sorted cells were added to anti-CD3- and anti-CD28-coated coverslips to assess the colocalization of α-Gal-3 IB with Gal-3, Alix, or ZAP-70. A and C. Representative confocal micrographs show the expression of Gal-3 IB or vector control, along with Gal-3, ZAP-70, or Alix, in stimulated CD8^+^ T cells. A 10 μm scale bar is shown. B and D. Bar graphs show cumulative data from the colocalization experiment, which is shown as Pearson’s colocalization coefficient (r) values. The levels of statistical significance were measured using One-way ANOVA and are shown as (****; p < 0.0001, ***; p < 0.001, **; p < 0.01, *; p ≤ 0.05, and ^ns^; p>0.05.

### α-Gal-3 IB-expressing CD8^+^ T cells exhibit an altered phenotype

Does the modified Gal-3 response by α-Gal-3 IB influence the activation and proliferative potential of stimulated CD8^+^ T cells? MACS-purified CD8^+^ T cells from lymphoid organs of WT mice were spin-transduced with control or the α-Gal-3 IB encoding pseudoviruses and stimulated with the plate-bound α-CD3/CD28 antibodies (Fig 3A and Fig. S3). The stimulated CD8^+^ T cells from Gal-3 KO were also included in the analysis to discern differences in the response patterns of cells modified by Gal-3 via genetic ablation or α-Gal-3-IB neutralization (Fig. 3A and Fig. S3). The change in the normalized values within each group was calculated, and differences amongst the Gal-3KO and α-Gal-3 IB groups were plotted as fold changes. Compared with unstimulated cells, both α-Gal-3 IB expressors and cells from Gal3-KO mice showed significantly higher expression levels of the early (CD69) and late (CD44) activation markers (Fig. 3B-D). While the α-Gal-3 IB expressing and Gal-3KO cells showed indistinguishable expression levels of CD69 at 24 and 48 hrs post stimulation, the former cell types showed up to 1.59-fold higher expression levels at 72 hrs (Fig. 3B and C, Fig 3D, upper panel). Similarly, the frequencies of CD44^+ve^ cells in the α-Gal-3 IB expressors as compared to the Gal-3 KO CD8^+^ T cells were also increased by 1.25 and 1.37-fold at 48 and 72 hrs of stimulation, respectively (Fig. 3B and C, Fig 3D, lower panel). The MFI values for CD69 and CD44 were also higher in α-Gal-3 IB-expressing cells as compared to the Gal-3 KO CD8^+^ T cells. (Fig. S3A and B). Therefore, α-Gal-3 IB, by modulating intracellular levels of Gal-3, induced a protracted activation response in the stimulated CD8^+^ T cells.

**Fig 3.**
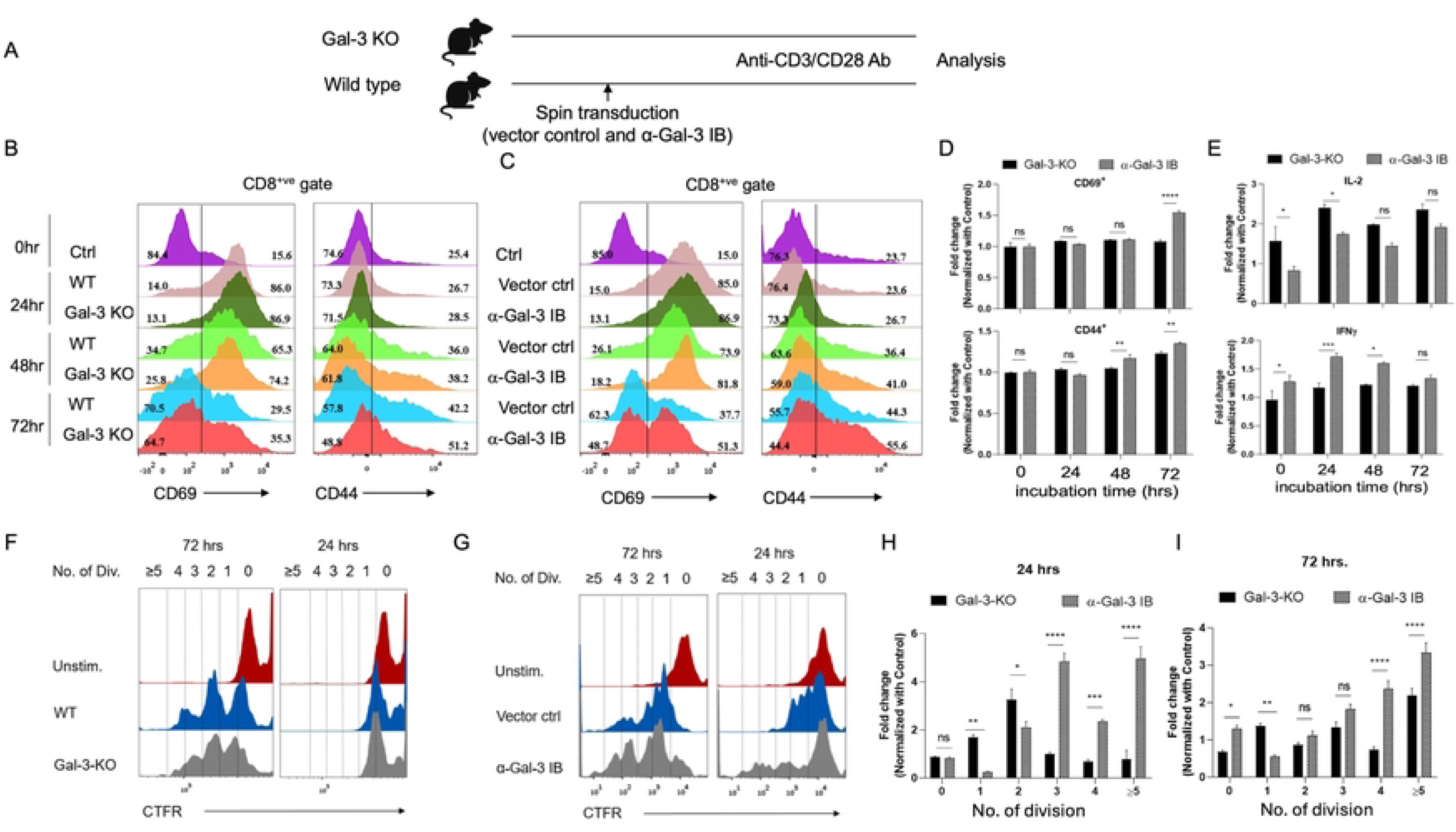
Assessing the *in vitro* responsiveness of Gal-3KO and α-Gal-3 IB expressing OT1 cells. A. CD8^+^ T cells from WT and Gal-3KO mice or the control transductants and α-Gal-3 IB expressors OT1 cells were stimulated with 1μg/ml of anti-CD3 and anti-CD28 antibodies. The expression of CD44 and CD69 was analyzed at different time points. B-C. The offset histograms are shown. The vertical line separates the CD44^hi^ and CD69**^+^**^ve^ cells from Gal-3 KO and WT mice (B) and the control transductants and the anti-Gal-3 IB expressing OT1 cells (C) at the indicated time post stimulation. D. Bar graphs show Mean ± SD values of the cumulative data with fold change values in anti-Gal-3 IB expressors compared to the cells from Gal-3 KO, based on the percent positive cells for the expression of CD69 and CD44. The fold change values in the Gal-3KO and α-Gal-3 IB groups were calculated by dividing the obtained values by those of the control groups (WT and VC, respectively). Two-way ANOVA was used for calculating statistical differences (****; p<0.0001, ***; p<0.001, **; p<0.01, *; p≤0.05 and ^ns^; p>0.05). E. At different time points during stimulation with anti-CD3 and anti-CD28, culture supernatants were collected to measure IL-2 and IFNγ concentrations released by the activated cells by ELISA. Bar graphs show the concentrations (Mean ± SD values) of released cytokines (pg/ml). The concentration of these cytokines was compared between different groups using Two-Way ANOVA (****; p < 0.0001, ***; p < 0.001, **; p < 0.01, *; p ≤ 0.05, and ^ns^; p>0.05). F-I. The naïve CD8^+^ T cells spin-transduced with VC and α-Gal-3 IB labeled with Cell Trace (Far Red) were stimulated with anti-CD3 and anti-CD28 antibodies at different time points to evaluate their proliferation potential. F-G. Representative offset plots showed the extent of proliferation in the WT and Gal-3 KO CD8^+^ T cells, as well as in VC and α-Gal-3 IB-expressing cells, along with their unstimulated controls at different time points. The dotted vertical lines were used to mark dividing cells. H-I. The cell frequencies in each division are shown in bar graphs. The values were plotted as Mean ± SD, and the experiments were performed 3-4 times. Two-way ANOVA was used to assess statistical differences and is shown as ****; p < 0.0001, ***; p < 0.001, **; p < 0.01, *; p ≤ 0.05, and ^ns^; p > 0.05.

Culture supernatants from the stimulated cells of different groups were collected to quantify cytokine levels. Gal-3 KO cells compared with those expressing α-Gal-3 IB produced higher levels of IL-2 at 24 hrs post-stimulation (Fig. 3E, upper panel). Furthermore, significantly higher levels of IFNγ were observed at 24 (1.82-fold) and 48 (1.63-fold) hrs in α-Gal-3 IB expressors (Fig. 3E). Therefore, α-Gal-3 IB-modified Gal-3 response in comparison to that achieved by a genetic disruption induced an elevated production of cytokines by the stimulated cells.

We also measured the proliferative potential of α-Gal-3 IB-expressing and Gal-3 KO CD8^+^ T cells. The GFP-positive cells were sorted and labeled with Cell Trace Far Red (CTFR). The labeled cells were then stimulated with plate-bound anti-CD3 and anti-CD28 antibodies to evaluate their proliferation potential (Fig. 3F-I and Fig. S3C and D). The α-Gal-3 IB expressors, compared with Gal-3 KO CD8^+^ T cells, exhibited an early proliferative response, with the former cell types showing several divisions within 24 hrs of stimulation (Fig. 3F-I). Furthermore, α-Gal-3 IB expressors, compared with Gal-3 KO CD8^+^ T cells, extensively proliferated both at 48 (5-fold) and 72 (4-fold) hrs (Fig. 3F-I and Fig. S3C and D). Therefore, modifying the Gal-3 response with IB, compared with the genetic knockdown approach, not only heightened the cytokine response but also enhanced the proliferation of the responding CD8^+^ T cells.

### Cell-intrinsic effects of Gal-3 in regulating the activation program of CD8^+^ T cells

Having established the improved response pattern of α-Gal-3 IB expressors, compared with Gal-3 KO cells, upon stimulation via their co-receptors, we assessed their antigen-specific activation profile. CTFR-labeled control or the α-Gal-3 IB expressing clonotypic T cells from OT1 TCR transgenic mice were co-cultured with Ovalbumin (Ova) or SIINFEKL peptide pulsed BMDCs from Gal-3 KO mice or those from WT mice with and without α-Gal-3 IB expression (Fig. 4 and Fig. S4). Furthermore, these experiments also allowed us to assess the contribution of Gal-3 produced by antigen-presenting cells (APCs) during T cell activation. BMDCs expressing α-Gal-3 IB and those from Gal-3 KO mice were first phenotypically analyzed (Fig. S4A-F). Compared with WT BMDCs, cells modified for a Gal-3 response, either by gene knockout or by α-Gal-3 IB expression, showed significantly higher levels of the costimulatory molecule CD86 (Fig. S4A-F). Thus, the frequencies of CD86^+ve^ cells were 41.3 ± 1.38% vs 51.2 ± 0.55% amongst Gal-3 KO and α-Gal-3 IB expressing BMDCs, respectively (Fig. S4A-F). The frequencies of cells expressing class I MHC (58.3 ± 0.27% vs 56.03 ± 0.95%) and class II MHC (59.8 ± 0.57% vs 57.86 ± 0.50%) were similar in Gal-3 KO and α-Gal-3 IB expressing BMDCs, respectively (Fig. S4A-F).

**Fig 4.**
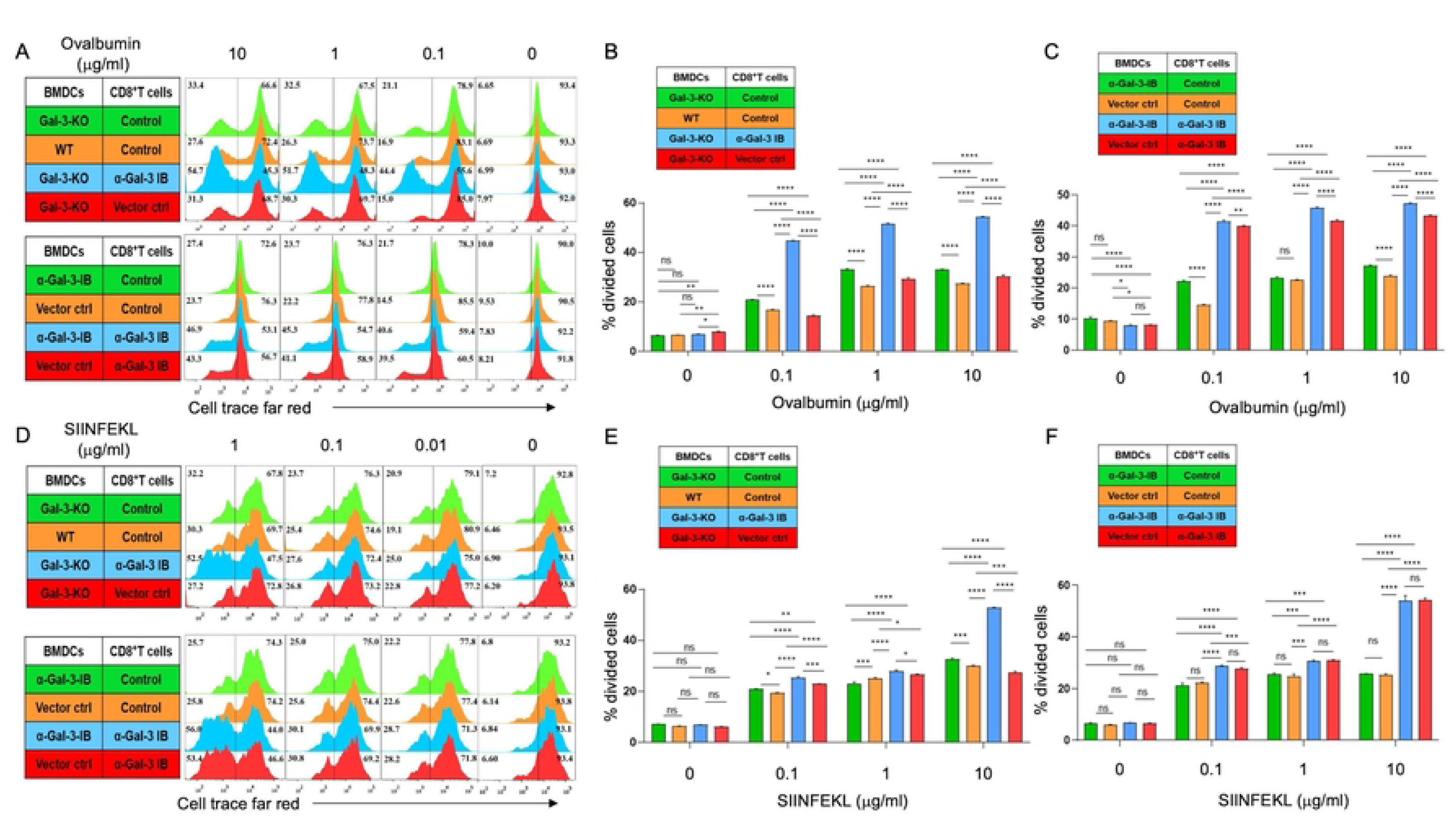
Assessing the cell-intrinsic effects of Gal-3 in influencing the proliferative response of CD8^+^ T cells stimulated by BMDCs via antigen presentation. A-F. BMDCs generated from bone marrow cells of WT and Gal-3KO mice or the WT BMDCs expressing α-Gal-3 IB or vector control were used for stimulating FACS-sorted OT1 cells. BMDCs were either stimulated with different concentrations of ovalbumin (0-10μg/ml) for 2 hours or pulsed with SIINFEKL peptide (0-1μg/ml) for 45 minutes. The pulsed BMDCs were co-cultured with Cell Trace Far-Red labeled α-Gal-3 IB, control, or the un-transduced OT1 cells. Proliferation of OT1 cells was measured. A and D. The offset histograms show the frequencies of proliferating cells with different division rates in the indicated conditions. B, C, E, and F. The frequencies of the dividing cells were plotted as bar graphs in the indicated conditions. Mean ± SD values are shown from one of the representative experiments. The experiments were performed three times. Two-way ANOVA was used to calculate statistical differences and is shown as (****; p < 0.0001, ***; p < 0.001, **; p < 0.01, *; p ≤ 0.05, and ^ns^; p>0.05.

The FACS-sorted BMDCs were then pulsed with the graded concentrations of Ova (0-10μg/ml) for 2 hrs and subsequently co-cultured with CTFR-labeled α-Gal-3 IB, vector transduced control cells, or the unmanipulated OT1 cells (Fig. S4G and H). The Ova-pulsed BMDCs of Gal-3 KO compared to those of WT mice induced slightly increased proliferation of OT1 cells at all the concentrations of Ova, with the frequencies of divided cells at 10μg/ml concentration being

33.4 ± 0.38% and 27.6 ± 0.35%, respectively (Fig. 4A and B). The α-Gal-3 IB expressing OT1 cells showed extensive proliferation compared to the control cells and such a response pattern remained unaltered by Gal-3 response of the stimulating BMDCs (Fig. 4A, upper panel and Fig. 4B). The frequencies of proliferating T cells were 54.7 ± 0.23% in the co-cultures of Gal-3 KO BMDCs and the α-Gal-3 IB OT1 cells (Fig. 4A, upper panel and Fig. 4B). Increased proliferation of α-Gal-3 IB expressing OT1 cells was also evident when such cells were co-cultured with control or the α-Gal-3 IB expressing Ova pulsed BMDCs with the frequencies of proliferating cells being 43.3 ± 0.21% and 47 ± 0.46%, respectively (Fig. 4A, lower panel and Fig. 4C). OT1 cells with the intact Gal-3 response showed reduced proliferation with the dividing cells being ∼23 ± 0.22% (Fig. 4A, lower panel and Fig. 4C). These results suggested that the Gal-3 response intrinsic to CD8^+^ T cells primarily regulated proliferation potential. Such effects were also evident at lower concentrations (1 and 0.1 µg/ml) of ovalbumin, with subdued effects observed at the lower concentration (Fig. 4A-C).

Whether or not the surface loading of peptide onto class I molecules in control or the α-Gal-3 IB-expressing BMDCs alters the activation of antigen-specific T cells was assessed. BMDCs from WT mice, Gal-3 KO mice, and the α-Gal-3 IB expressing WT BMDCs were pulsed with varying concentrations of SIINFEKL peptide (0.01-1μg/ml) for 45 minutes and subsequently co-cultured with the CTFR-labeled control or the α-Gal-3 IB expressing OT1 cells. As compared to the control cells, the OT1 cells expressing α-Gal-3 IB showed more than 2-fold higher frequencies (56.5 ± 0.13% vs 27.1 ± 0.35%) at 1μg/ml of the peptide concentration (Fig. 4D, upper panel and Fig. 4E). Furthermore, the BMDCs modified of Gal-3 response either by gene knockout or using the α-Gal-3 IB did not significantly alter the proliferative response of T cells (Fig. 4D, lower panel and Fig. 4F). These experiments demonstrated that irrespective of the nature of stimulating BMDCs, the α-Gal-3 IB-expressing CD8^+^ T cells responded more efficiently to antigenic response compared to control cells.

### α-Gal-3 IB-expressing CD8^+^ T cells efficiently expand during virus infection

α-Gal-3 IB-expressing CD8^+^ T cells compared with their control counterparts generated efficient activation and proliferative responses *in vitro*. We then assessed their responsiveness during the resolution of influenza A virus infection. 40,000 of FACS-purified control and α-Gal-3 IB expressing OT1 cells (CD45.2^+^) were transferred into sex-matched congenic CD45.1^+^ mice, which were then infected intranasally with 200pfu of Influenza A virus encoding SIINFEKL, an H-2K^b^ restricted epitope (IAV: WSN-SIINFEKL) (Fig. 5A and S5A-D). Control and α-Gal-3 IB-expressing cells showed similar survivability, with the proportion of dead cells (propidium staining (PI)-positive) being 12.37 ± 0.68% and 13.7 ± 0.56%, respectively (Fig. S5A-D). The frequencies of donor cells were measured in peripheral blood before the primary infection to ensure equal transfer (Fig. S5E). The response pattern of the donor cells was then analyzed during the acute phase by 7 days post-infection (dpi) and in the early memory phase at 45 dpi in the lymphoid and non-lymphoid organs. The recall response of the persisting memory cells was measured three months later by infecting the animals with a heterologous gamma herpesvirus encoding SIINFEKL epitope (MHV-68-SIINFEKL) (Fig. 5A and S6). In the acute phase of the response, the frequencies of virus-specific donor CD8^+^ T cells (CD45.2^+^CD44^+^) were 1.5 to 2-fold higher in the peripheral blood of animals receiving α-Gal-3 IB expressing CD8^+^ T cells as compared to those receiving the control cells (Fig. 5B and C). The α-Gal-3 IB CD8^+^ T cells compared to the control cells at 3dpi, 5dpi, and 7dpi were 2.2 ± 0.09% vs 0.88 ± 0.25%, 3.2 ± 0.16% vs 1.99 ± 0.80%, and 5.36 ± 0.95% vs 3.58 ± 0.43%, respectively (Fig. 5B and C). Interestingly, the frequencies of responding donor cells (CD44^+^CD45.2^+^) remained 2- to 3-fold higher in recipients of the α-Gal-3 IB CD8^+^ T cells than those receiving the control cells until 45 dpi (Fig. 5B and C). The recallability of α-Gal-3-IB-expressing CD8^+^ T cells and vector-transduced cells was assessed at 7 days post-recall (dpr). The frequencies of donor cells were 2.69 ± 0.05% and 1.64 ± 0.05% in the recipients of α-Gal-3 IB OT1 cells, as compared to those receiving control OT1 cells (Fig. 5B and C).

**Fig 5.**
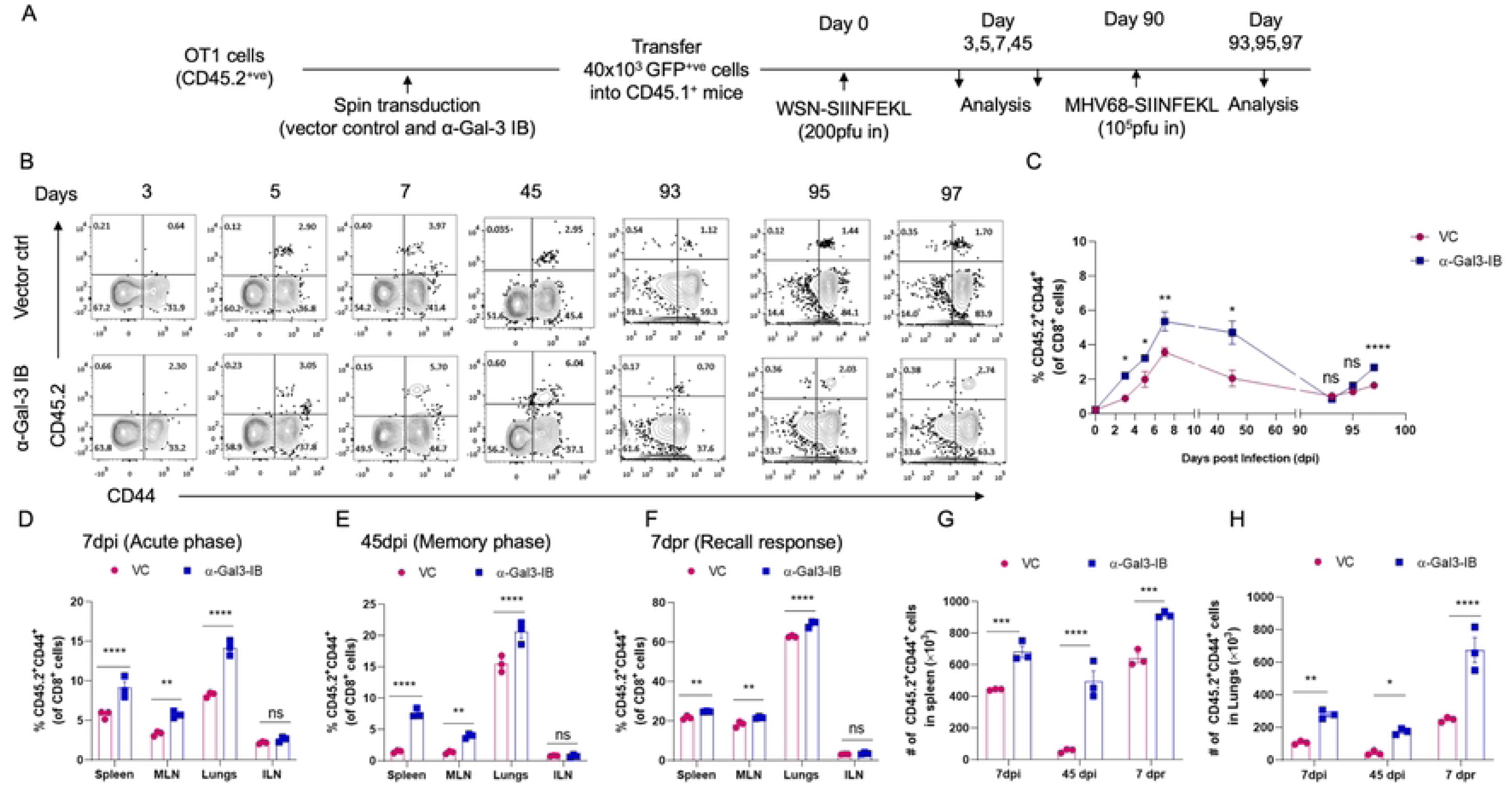
Prospective analysis of the expanded antigen-specific control and α-Gal-3 IB expressing OT 1 cells during the primary and the memory response. A. The frequencies of the donor cells were measured in peripheral blood and different organs in the primary (7dpi) and memory (45dpi) response, as well as at different times post-rechallenge of mice with MHV68-SIINFEKL. B. Representative FACs plots show the frequencies of virus-specific donor (CD45.2^+^CD44^+^) cells in the peripheral blood of mice receiving vector control or the α-Gal-3 IB expressing OT1 cells at the indicated time points. C-F. The frequencies of virus-specific donor cells (CD45.2^+^CD44^+^ or CD45.2^+^) in each group are depicted by line diagrams at the indicated time points in peripheral blood (C), different lymphoid organs, as well as lung tissues at 7dpi (D), 45dpi (E), and 7 days post recall (F). G-H. Absolute counts of donor cells are shown at different time points in the spleen (G) and lungs (H). Mean ± SD are shown, and the level of statistical analysis is determined by Two-way ANOVA and is shown as (****; p < 0.0001, ***; p < 0.001, **; p < 0.01, *; p ≤ 0.05, and ^ns^; p>0.05.

We also measured the frequency and absolute number of antigen-specific donor cells in both lymphoid organs and lungs of animals during primary as well as secondary infections (Fig. 5D-H). During the primary response, the frequencies of donor cells in the recipients of α-Gal-3 IB CD8^+^ T cells as compared to those receiving control cells were 9.11 ± 1.35% vs. 5.7 ± 0.50% in spleen, 5.72 ± 0.45% vs. 3.32 ± 0.30% in draining mediastinal lymph node (Med LN) and 14.1 ± 0.95% vs. 8.2 ± 0.33% in lung tissues at 7dpi (Fig. 5D-H and Fig. S6B). At 45 dpi, the frequency of the CD44^+^CD45.2^+^ in the recipients of α-Gal-3 IB CD8^+^ T cells, as compared to those receiving control cells, was 7.59 ± 0.73% vs 1.47 ± 0.21% in spleen, 3.96 ± 0.39% vs 1.34 ± 0.22% in Med LN, and 20.60 ± 1.80% vs 15.46 ± 1.30% in lungs (Fig. 5D-E and Fig. S6B). While the antigen-specific CD8^+^ donor T cells were detectable in non-draining lymph nodes, no significant differences were observed between the control and the α-Gal-3 IB expressors (Fig. 5D-E). Thus, the α-Gal-3 IB expressors were present in 2 to 4-fold excess compared to the control cells in different organs except for the non-draining inguinal LNs. These results suggest an efficient expansion of α-Gal-3 IB expressors compared to the control cells, which could improve their memory transition.

The recall response of the memory cells was measured in different lymphoid and non-lymphoid organs. The recipients of α-Gal-3 IB CD8^+^ T cells exhibited a significantly improved recall response compared with those receiving the control cells (Fig. 5 and Fig. S6). The frequencies of CD44^+^CD45.2^+^ cells in the recipients of α-Gal-3 IB CD8^+^ T cells and the control cells were 24.76 ± 0.21 vs 21.66 ± 0.96% in spleen, 21.73 ± 0.51% vs 18.33 ± 1.48% in Med LN, and 69.2 ± 1.74% vs 62.8 ± 0.52% in lungs, respectively (Fig. 5F and S6 B). No differences were observed in the frequencies of CD44^+^CD45.2^+^ cells in the non-draining inguinal LNs of the two groups. The absolute numbers of donor CD44^+^CD45.2^+^ cells, however, were 1.5- to 3-fold higher in different lymphoid organs and lungs of animals receiving α-Gal-3 IB CD8^+^ T cells compared to those of the control cells (Fig. 5G-H and Fig. S6C-D).

### α-Gal-3 IB-expressing CD8^+^ T cells exhibit superior functionality

We compared the functionality of α-Gal-3 IB and control cells recruited in the response to IAV infection by measuring their cytokine production using an intracellular cytokine staining (ICCS) assay following *in vitro* stimulation with SIINFEKL peptide (Fig 6 and Fig S7). The frequencies of IFNγ^+^, TNFα^+^, and IFNγ^+^TNFα^+^ double-positive (DP) cells across different organs were assessed in both the primary and memory response. A significant increase in the frequencies of IFNγ^+^ α-Gal-3 IB CD8^+^ T cells compared to the vector control cells was observed in spleen (45.07 ± 1.46% vs 37.74 ± 3.07%), MedLN (26.39 ± 3.33% vs 14.65 ± 1.93%), and lungs (29.43 ± 1.05% vs 23.72 ± 2.22 %) at 7dpi (Fig. 6 A and Fig. S7B). Similarly, the α-Gal-3 IB expressing CD8^+^ T cells compared to the control cells had higher frequencies of IFNγ^+^ TNFα^+^ DP cells in spleen (15.3 ± 0.87% vs 11.36 ± 1.20%) and lungs (8.567 ± 1.37% vs 5.2 ±1.76%) (Fig. 6 B-C, Fig. S7B).

**Fig 6.**
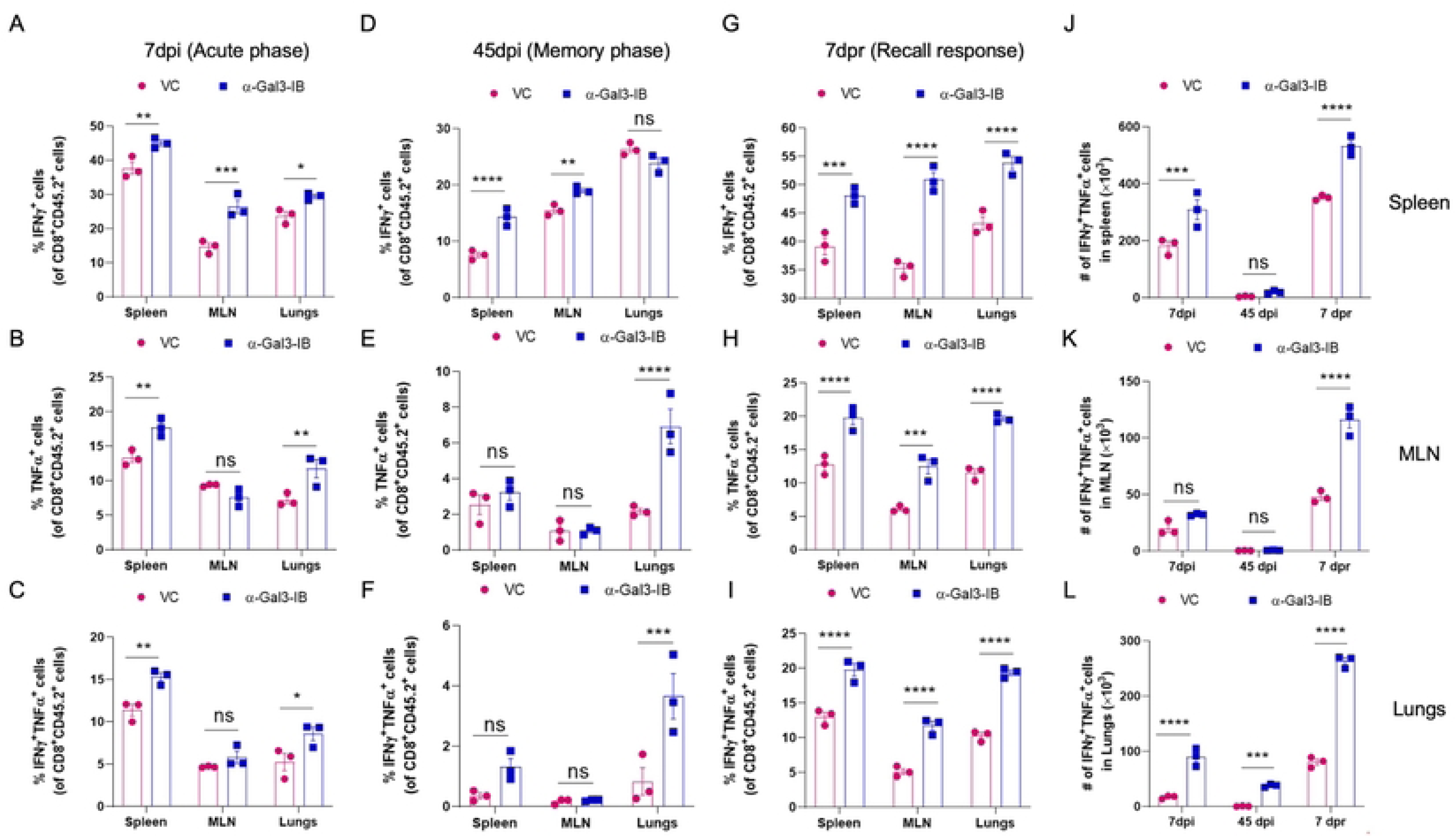
Assessing the functionality of antigen-specific α-Gal-3 IB expressors and control CD8^+^ T cells. The functionality of the virus-specific vector-transduced, and α-Gal-3 IB expressing CD8^+^ T cells was assessed in different lymphoid organs and lungs of mice infected with WSN-SIINFEKL IAV in the acute (A-C), memory (D-F), and the recall phase (G-I). The prepared single cell suspensions were stimulated with SIINFEKL peptide for 4 hours at 37°C in the presence of Brefeldin A. The cells were first stained for cell-surface markers, then permeabilized to stain for the accumulated cytokines using a standardized procedure for ICCS assays. Frequencies and cell numbers in the different organs were normalized to those of unstimulated cells. A-C. The frequencies of IFNγ^+^, TNFα^+^, and IFNγ^+^TNFα^+^ double-positive (DP) cells in different organs were quantified and shown with bar graphs at 7dpi (A-C), 45dpi (D-F), and 7days post recall (dpr) of memory response at three months (G-I). Absolute counts of the IFNγ^+^ TNFα^+^ DP cells are shown by bar graphs at different time points in the spleen (J), Med LN (K), and lungs (L). Mean ± SD values for the frequencies and numbers of cytokine-producing cells are shown. Two-way ANOVA was used to calculate statistical differences and is shown as (****; p < 0.0001, ***; p < 0.001, **; p < 0.01, *; p ≤ 0.05, and ^ns^; p>0.05.

In the memory response at 45 dpi, the α-Gal-3 IB expressing CD8^+^ T cells, as compared to the control cells, showed ∼3-fold higher frequencies of IFNγ^+^TNFα^+^ (3.65 ± 1.30% vs 0.83 ± 0.80%) in lung tissues (Fig. 6D-F, Fig. S7B). The recalled α-Gal-3 IB expressors compared to the control cells also showed a significantly enhanced frequencies (Fig. 6G-I and Fig. S7B-E) and absolute numbers (Fig. 6J-L and Fig. S7B-E) of both IFNγ^+^ as well as TNFα^+^ cells isolated from lymphoid organs and lung tissues, both in terms of frequency (more than 2-fold) and absolute numbers (upto 3-fold) after three months of primary infection. Taken together, the α-Gal-3 IB expressors expanded following IAV infection and generated a more efficient cytokine response than the control cells.

### The α-Gal-3 IB expressing CD8^+^ T cells expanded following IAV infection preferentially generate memory precursor effector cells (MPECs)

Since the α-Gal-3 IB expressors elicited a stronger memory response than the control cells, we assessed whether this programming is determined early in their differentiation. The expression of IL-7 receptor (CD127) and killer cell lectin-like receptor G1 (KLRG1) in the expanded cells during the acute phase of a response can help identify memory precursor effector cells, MPECs (CD127^+^ KLRG1^-^) or the short-lived effector cells, SLECs (CD127^-^KLRG-1^+^) phenotype (26). While the SLECs help resolve the infection by cytolyzing infected cells, MPECs further differentiate into persisting memory cells (27). The proportions of MPECs among the donor α-Gal-3 IB-expressing cells and the control cells were 34.33 ± 2.07 % vs 14.2 ± 0.44 % in spleen, 26.4 ± 1.80% vs 17.43 ± 1.02% in the Med LN, and 16 ± 1.10 % and 9.08 ± 1.76 % in lungs of the IAV-infected animals at 7dpi (Fig. 7A and B). On the contrary, the frequencies of SLECs were significantly lower within the α-Gal-3 IB expressors as compared to the control cells in spleen (12.3 ± 0.95% vs 22.63 ± 0.50%), Med LN (5.86 ± 0.43 % vs 9.53 ± 0.481%), and lungs (34.1 ± 3.09% vs 55.46 ± 0.45%), respectively (Fig. 7A and C). These results suggested that the inhibited Gal-3 response by α-Gal-3 IB in CD8^+^ T cells early during the activation process preferentially induced MPECs over SLECs.

**Fig 7.**
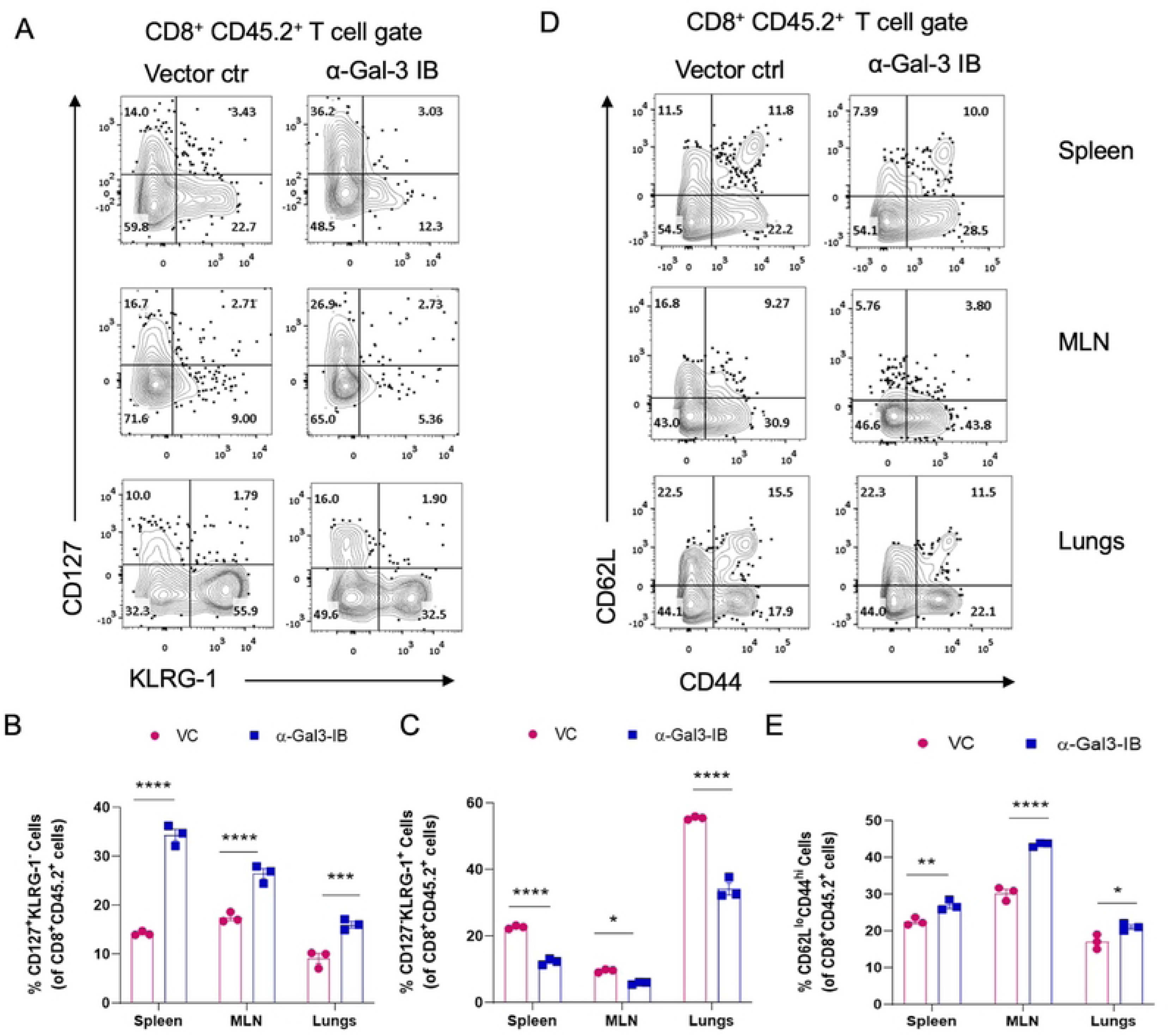
The α-Gal-3 IB CD8^+^ T cells preferentially generate MPECs over SLECs. The vector-transduced cells and the α-Gal-3 IB expressors that expanded during IAV infection were phenotypically characterized at 7dpi. The proportion of SLECs and MPECs in both groups was assessed by staining the cells with anti-KLRG1 and anti-CD127 antibodies. A. Representative FACS plots show the frequencies of CD127^+^KLRG-1^-^ (MPECs) or CD127^-^KLRG-1^+^ (SLECs) cells in spleen, Med LN, and lungs of infected animals at 7dpi. Bar graphs show the proportion of MPECs (B) and SLECs (C) in different organs. D. Representative FACS plots show the frequencies of activated cells (CD62L^lo^CD44^hi^) in both groups. E. Frequencies of CD62L^lo^CD44^hi^ cells are plotted as bar graphs. Two-way ANOVA was used to calculate the levels of statistical significance, which are indicated as ****; p < 0.0001, ***; p < 0.001, **; p < 0.01, *; p ≤ 0.05, and ^ns^; p >0.05.

Phenotypic analysis of the donor cells at 7dpi further revealed that the α-Gal-3 IB expressors, as compared to the vector control cells, had significantly higher frequencies of CD62L^lo^CD44^hi^ cells, with their proportions being 26.93 ± 1.40% vs 22.5 ± 1.04% in spleen, 43.5 ± 0.61% vs 30.23 ± 1.89% in Med LN and 21.03 ± 1.05% vs 17.0 ± 2.0% in lungs, respectively (Fig. 7D and E). These results suggested that the α-Gal-3 IB-expressing cells exhibited a more activated phenotype than the control cells (Fig. 7D and E).

### α-Gal-3 IB-expressing CD8^+^ T cells preferentially generate effector memory (T_EM_) cells

We assessed the phenotypic attributes of the differentiated antigen-specific CD8^+^ T cells generated from the control and α-Gal-3 IB-expressing cells during the memory phase at 45 dpi. The proportions of CD62L^lo^CD44^hi^ (a phenotype commonly attributed to the effector memory cells; T_EM_) amongst the α-Gal-3 IB expressors compared to the control cells were 34.63 ± 2.52% vs 27.3 ± 0.95% in spleen, 36.2 ± 1.83% vs 30.73 ± 1.51% in Med LN, and 61.87 ± 3.01% vs 56.56 ± 1.20 % in lungs, respectively (Fig. 8A and B). On the contrary, CD62L^hi^CD44^hi^ cells (central memory cells; T_CM_) amongst the α-Gal-3 IB cells and the vector cells were 47.4 ± 0.60% vs 52.2 ± 1.20% in spleen, 52.8 ± 2.05% vs 58.46 ± 1.00% in Med LN, and 28.8 ± 0.61% vs 39.6 ± 1.15% in lungs, respectively (Fig. 8A and C). We also observed that the α-Gal-3 IB expressors, as compared to the control cells, showed decreased frequencies of CCR7^+^ cells, with these cells being 31.8 ± 0.95% vs 43.6 ± 0.34% in splenic tissues, 53.3 ± 1.67% vs 59.5 ± 1.31% in Med LN, and 21.93 ± 0.51% vs 33.26 ± 3.04% in lungs, respectively (Fig 8D and E). CCR7 facilitates the homing of CD8^+^ T cells to secondary lymphoid organs (28,29). The MFI values of CCR7 were 504.3 ± 7.10% vs 650 ± 51.68% in splenic tissues, 741.66 ± 16.04% vs 825.66 ± 37.86% in Med LN, and 489 ± 12.0% vs 590 ± 14.29% in lungs, respectively (Fig. 8F). The lower expression of CCR7 by antigen-specific cells also indicates a TEM phenotype (30,31). Taken together, the phenotypic analysis of differentiating α-Gal-3 IB expressors and control cells suggested that anti-Gal-3 IB preferentially induced a TEM phenotype by modifying the Gal-3 response.

**Fig 8.**
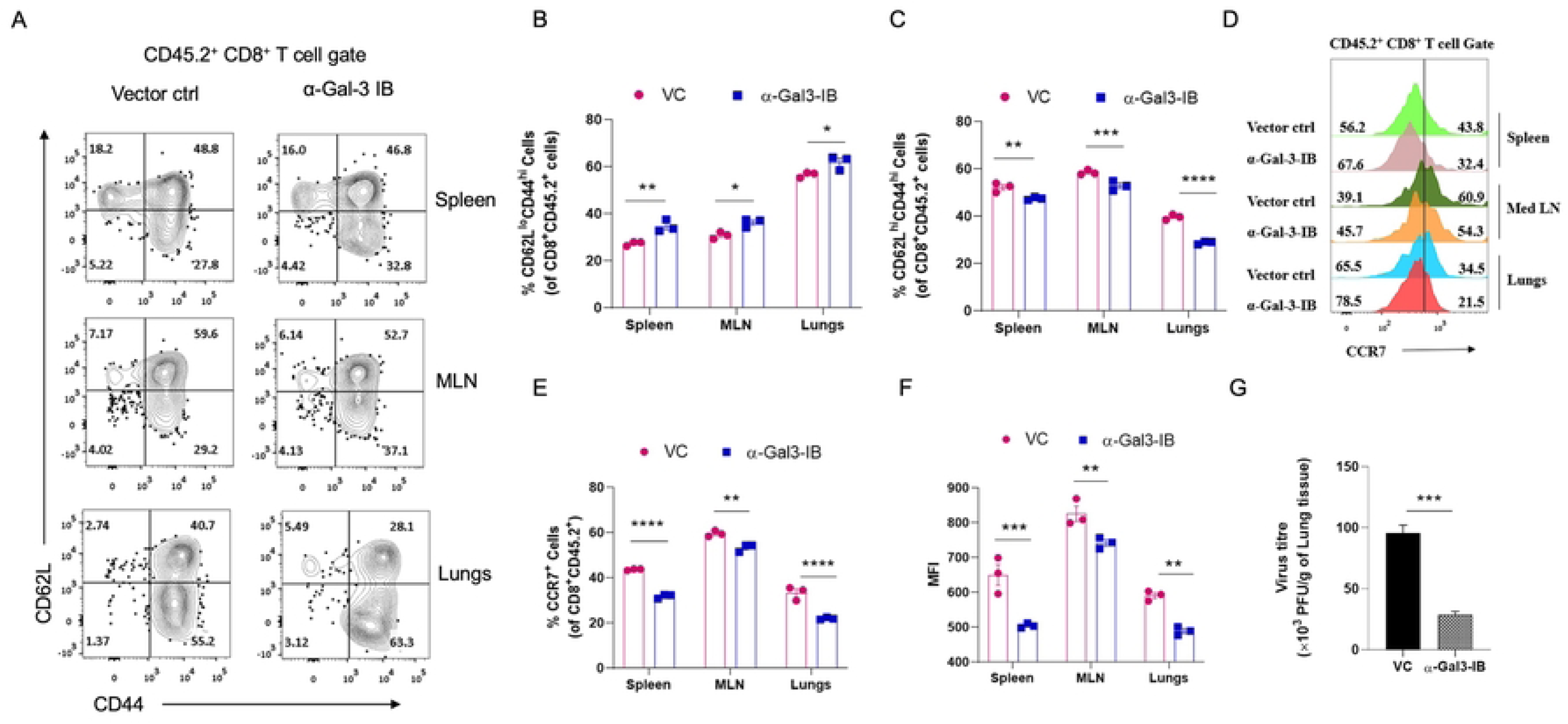
The α-Gal-3 IB expressors that respond to IAV infection preferentially generate T_EM_ phenotype and control infection better upon recall. The vector-transduced and anti-Gal-3 IB-expressing cells were phenotypically characterized at the memory stage at 45dpi. Frequencies of CD62L^lo^CD44^hi^ and CD62L^hi^CD44^lo^ cells were analyzed during the memory phase, as shown by representative FACs plots (A) and by bar graphs (B and C). Frequencies of CCR7^+^CD8^+^ T cells amongst both the vector transduced and the anti-Gal-3 IB expressing cells are shown by representative offset histograms (D) and bar graphs (E). F. MFI values for the expression of CCR7 are shown by bar graphs. Mean ± SD values are shown, and a Two-way ANOVA was used to calculate statistical differences. G. The animals infected with WSN-SIINFEKL IAV were re-infected after 90 days with a γ-herpes virus (MHV-68-SIINFEKL), and the viral loads were determined in the lung tissues at 7 days post-secondary infection by plaque assays. Mean ± SD values are shown by bar graphs. An unpaired t-test was used to calculate the statistical difference between the two groups. ****; p < 0.0001, ***; p < 0.001, **; p < 0.01, *; p ≤ 0.05, and ^ns^; p>0.05.

We also compared viral loads in animals re-infected with MHV68-SIINFEKL. We observed up to a five-fold reduction in MHV68 titers in the groups receiving α-Gal-3 IB-expressing cells compared with the group receiving vector-transduced cells. Therefore, the α-Gal-3 IB expressing cells helped control the infection better (Fig. 8G).

To establish whether the intracellular expression of α-Gal-3 IB persists in the α-Gal-3 IB expressors, we quantified the number of c-Myc^+ve^ cells as the IB was expressed as a c-Myc tagged protein. After seven days of recall response, 35.53 ± 1.76 % in spleen, 31.13 ± 1.63 % in Med LN, and 50.7 ± 1.57% in lungs of the expanded CD8^+^ T cells scored positively with an anti-c-Myc antibody (Fig. S8A and B). The positivity for c-Myc staining was evident amongst the α-Gal-3 IB expressors in absolute numbers as well as the MFI values, while the vector control cells showed only a minimal level of background staining (Fig S8C and D).

## Discussion

We compared the differentiation pattern of antigen-specific CD8^+^ T cells that were depleted of Gal-3 response using an α-Gal-3 IB during a resolving respiratory infection caused by Influenza A virus. The α-Gal-3 IB-expressing cells showed an elevated and sustained activation, enhanced proliferation, and increased cytokine production compared to the genetically depleted cells from Gal-3 KO mice. The α-Gal-3 IB expressing clonotypic OT1 cells generated an efficient effector response with skewed ratios of MPECs (CD127^hi^KLRG1^lo^) over SLECs (CD127^lo^KLRG1^hi^), leading to an efficient effector memory formation following infection with a resolving Influenza A virus encoding Ovalbumin-derived SIINFEKL epitope. The recalled α-Gal-3 IB-expressing cells more effectively controlled the secondary infection. We argue that IB-mediated modulation of protein-protein interactions in CD8^+^ T cells is a viable approach to fine-tune T cell differentiation program to achieve a favorable outcome of infection. Such a strategy could be valuable in managing T cell-mediated immunopathological conditions, as well as cancers of different types (32).

Gal-3, due to its high affinity for glycans, regulates the maturation and differentiation programs of T cells during developmental processes, modulates their migratory patterns and survival during thymocyte-stromal cell interactions. Subsequently, Gal-3 influences the activation and differentiation of T cells in the periphery, thereby affecting the balance between immunity and tolerance (33,34). Many investigators have attempted to elucidate the inhibitory role of Gal-3 in CD8^+^ T cell differentiation by studying its interactions with CD45, LAG-3, LAT, CTLA-4, LFA-1, and MGAT-5 (11,35–37). Several studies have shown that complete genetic ablation of Gal-3 in CD8^+^ T cells, from Gal-3KO mice, improves their effector functions (12,38–40). Recently, it was reported that Gal-3 acts as a ‘physiological handbrake’ in regulating the effector functions of cytotoxic T cells. Accordingly, Gal-3-depleted T cells produced significantly elevated levels of effector cytokines and, in doing so, affected the allogeneic hematopoietic cell transplantation outcomes (41). The role of intracellular versus extracellular Gal-3 in the regulation of T cell differentiation has not been clearly defined. Gal-3, when expressed intracellularly, interacts with the Alix protein and downregulates the TCRs (25). Others and we have shown previously that Gal-3 is recruited to the immune synapse to regulate the activation of both naive and memory CD8^+^ T cells, and that the CD8^+^ T cells from Gal-3 KO mice responded more efficiently than those from WT mice during a γ-herpesvirus (MHV68) infection (12,25,42). Currently, there is limited data available to define the contribution of intracellular Gal-3 in the activation, differentiation, or migration of CD8^+^ T cells.

To establish gene function, genetic modification approaches such as siRNA or the knock-out techniques are employed. However, such procedures can pose risks of off-target effects, irreversible alterations in cellular development and physiology, and introduce genetic scars, which could all contribute to over or underestimation of the real effects (16). Small molecule inhibitors used to modify protein functions also show side effects, including commonly observed cytotoxic effects, as observed with some kinase inhibitors, which trigger serious cardiovascular complications (43,44). This necessitates devising alternative strategies. Conventional antibodies are designed to function in extracellular spaces. In contrast, V_H_H derivatives of camelid heavy-chain-only antibodies, also known as nanobodies or sdAbs, are small-sized binders, highly stable, and exhibit unique structural and functional features. These binders can be efficiently expressed intracellularly as functional products with intact antigen-binding capabilities (45). Therefore, intrabodies can disrupt macromolecular interactions by sequestering the available ligand and can help unmask the physiological effects of a specific protein (46). Given their expanding utility as recognition modules for chimeric antigen receptors (CARs) and for generating nanobody-based killer T cell engagers (TCEs), we explored the potential of V_H_H in modifying the intracellular functions of Gal-3 in CD8^+^ T cells (21,47). We used α-Gal-3 IB, which specifically interacted with the CRD of Gal-3, to modulate the bioavailable levels of Gal-3 in CD8^+^ T cells and assessed their activation and differentiation patterns during a resolving IAV infection (48). Furthermore, to minimize the effects of varying affinities of TCRs expressed by polyclonal T cells, we modulated the Gal-3 response in clonotypic OT1 cells using α-Gal-3 IB.

Following stimulation, both α-Gal-3 IB-expressing OT1 cells and Gal-3KO CD8^+^ T cells showed higher frequencies of CD69^+^ and CD44^+^ cells compared to their respective control groups. However, when the data were normalized to assess differences, the frequencies of cells expressing both markers in α-Gal-3-IB-expressing cells compared with Gal-3 KO cells were higher, particularly at later time points, indicating that α-Gal-3-IB-expressing cells exhibited a sustained activation profile (Fig. 3B-D). Such cells also produced higher levels of IL-2 and IFN-γ (Fig. 3E) and proliferated more extensively as compared to those genetically depleted of Gal-3 (Fig. 3F-I and Fig. 4). Gal-3 KO animals produced elevated levels of L-2 following *Trypanosoma cruzi* infection (49). The high concentration of IFN-γ in the supernatants of activated α-Gal-3 IB-expressing CD8^+^ T cells provided an early indication of their enhanced effector functions. Intracellular Gal-3 has a complex relationship with IFN-γ production. Accordingly, it was observed that the intracellular Gal-3 influenced IFNγ signaling (38,50,51). Elevated IFN-γ production could help control viral load more effectively. Thus, recipients of α-Gal-3 IB-expressing T cells, compared with those receiving control cells, efficiently controlled the rechallenge virus. The modification of the Gal-3 response, either by genetic depletion or by an α-Gal-3 IB in the stimulating APCs, did not significantly alter the T cell response pattern, suggesting that the Gal-3 produced by T cells primarily regulated their functionality (Fig. 4).

It has also been suggested that Gal-3 exerts immunomodulatory effects by decreasing TCR affinity for its cognate MHC-I peptide, thereby inducing efficient internalization of TCRs. This causes enhanced apoptosis of T cells as well as their inefficient stimulation (52–54). Inhibiting intracellular Gal-3 using α-Gal-3 IB in T cells, which otherwise develop in a normal microenvironment, and their analysis could provide direct evidence on the intracellular role of Gal-3. Thus, antigen-specific T cells expressing a clonotypic TCR were analyzed using an adoptive transfer approach to measure their differentiation in a Gal-3-sufficient microenvironment of WT mice. The α-Gal-3 IB-expressing cells exhibited enhanced effector functions and generated improved functional memory. They displayed a more activated (CD62^lo^CD44^hi^) phenotype than the control cells, suggesting that Gal-3 response actively regulates the T cell differentiation program (Fig. 5-8). Even after three months of persistence in the recipient animals, the α-Gal-3 IB-expressing cells consistently scored highly for c-Myc positivity, suggesting their active regulation of the Gal-3 response (Fig S8). While the molecular details are being analyzed in the Gal-3-mediated fine-tuning of T cell responses, more efficient TCR signaling in the responding cells following Gal-3 inhibition is evident.

We demonstrate that modulating Gal-3 response intracellularly in CD8^+^ T cells using α-Gal-3 single-domain antibodies can be a readily usable strategy to improve their effector function by seamlessly bypassing undesired, promiscuous off-target effects. The approach could have a broader applicability in deciphering intracellular molecular interactions and identifying novel targets that can help modulate cellular response. The α-Gal-3 IBs offer a viable option to enhance effector functions and overcome the immunosuppressive barriers not only to achieve favorable outcomes in infections but also during tumorigenesis.

## Materials and Methods

### Mice and viruses

The strain of mice viz., C57BL/6 (Stock No. 000664), B6 OT1 (*C57BL/6-Tg(TcraTcrb)1100Mjb/J*; Stock No. 003831), and B6 CD45.1 (*B6.SJL-Ptprca Pepcb/BoyJ*; Stock No. 002014) were procured from Jackson Laboratory, USA. The animals were housed and bred in the individual ventilated cages in the animal facility of Indian Institute of Science Education and Research (IISER), Mohali. The Institutional Animal Ethics Committee (IAEC), IISER Mohali, constituted under the aegis of the Committee for the Purpose of Control and Supervision of Experiments on Animals (CPCSEA) approved all the animal protocols and the experiments were performed accordingly. MHV68-SIINFEKL and Influenza A Virus (IAV, WSN-SIINFEKL) were used for the *in vivo* experiments. MHV68-SIINFEKL was propagated, harvested and titrated using Vero cells and stored at -80°C until further use. WSN-SIINFEKL was propagated and titrated using MDCK cells.

### Antibodies and other reagents

Antibodies used for measuring the expression of different molecules were procured from BD Biosciences, Tonbo biosciences, eBiosciences and BioLegend. The following antibodies were used CD8-PerCP-Cy5.5 (53-6.7), CD45.2-APC, -FITC & -PE (1O4), CXCR3-FITC (173), CD44-APC (IM7), CD62L-APC (MEL 14), CD69-APC (H1.2F3), CD127-APC, -FITC (A7R34), KLRG1-AF488 (2F1), CD103-FITC (2E7), IFN-γ-PE (XMG1.2), TNF-α-APC (TN3-19), Granzyme B-PE (NGZB), CCR7-APC (4B12), CD80-PE (16-10A1), CD86-APC (GL-1), MHCI-APC (28-8-6) MHCII-APC (M5/114.15.2). SIINFEKL (Ova257-264) peptide was used for stimulating cells in intracellular cytokine staining assays. All the antibodies were diluted in FACS buffer (Phosphate Buffered Saline (PBS) with 2% FBS) in 1:200 ratio. MojoSort CD8^+^ T cell purification kit was purchased from BioLegend. CFSE, intracellular fixation-permeabilization buffer, and Streptavidin-PE were purchased from Thermofisher. Monensin, Brefeldin A, purified α-CD3ε (17A2) and α-CD28 (37.51) were purchased from eBioscience.

### Cloning of α-Gal-3 sdAb in pLenti-GFP vector

Anti-Gal-3 sdAb was cloned downstream to the CMV promoter, between Xba-1 and BamH-1 sites of the mammalian expression vector; pLenti GFP. The α-Gal-3 sdAb was amplified using the sdAb-specific forward primer containing the *Xba1* site and the sdAb-specific reverse primer encoding the c-Myc tag with the *BamH1* site. After amplification, the amplicon was digested with the specific restriction enzymes, ligated into the already digested pLenti-GFP vector, and transformed into the Stbl3 strain of *E. coli*. The positive colonies were confirmed by colony PCR and restriction digestion. The positive clones were grown in the LB media for plasmid isolation.

### Preparation of α-Gal-3 sdAb-expressing pseudoviruses

Anti-Gal-3 IB was generated by transfecting 80% confluent HEK293 T cells with 15μg of the α-Gal-3 sdAb containing pLenti-GFP vector, 9.5μg of the packaging vector-pCMVR8.74, 15μg of the envelope vector-pMD2.G, and 5μg each of TAT and REV plasmids (24,55). For the vector control, only pLenti-GFP vector along with the other plasmids were used at same concentrations.

All these plasmids were added to serum-free DMEM containing PEI at a 1:3 ratio and the transfection mixture was vortexed for 20 seconds and then left in stationary phase for 15 minutes at RT. Thereafter the mix was added to HEK293T cells. Five-hours after transfection, the medium was replaced with 10% DMEM, and after 72 hours, the supernatants containing control or the a-Gal-3 IB encoding peudoviruses were collected and precipitated using a PEG-NaCl solution. After precipitation, the pellets were dissolved in 1ml serum-free DMEM and used to transduce CD8^+^ T cells or the BMDCs.

### Transduction of CD8^+^ T cells and BMDCs to express α-Gal-3 IB

Spleen and lymph nodes were pooled from either C57BL/6 or OT1 mice in 10% RPMI and subjected to RBC lysis after preparing the cell suspension. After RBC lysis, the cells were centrifuged at 1380 rpm for 15 minutes at 4 °C, and the pellet was dissolved in the media. CD8^+^T cells were MACS-purified using the provided protocol. After MACS, the purity of the CD8^+^T cells was checked, and they were divided into two groups: the inactivated or naïve CD8^+^T cells and the in vitro activated CD8^+^T cells, which were activated using anti-CD3 and anti-CD28 antibodies for 48 hours. These cells were then spin-transduced in 96-well U-bottom plates using pseudoviruses expressing anti-Gal-3 sdAb (i.e., IB) or the vector control (VC). The spin-transduction was carried out for 90 minutes at 800g at 37 °C. The next day, the media was replaced with 10% RPMI, and after 72 hours, GFP-positive CD8 T cells were analyzed and sorted for different experiments.

Bone marrows were collected from the long bones such as femur and tibia of euthanised C57BL/6 mice. The bones were flushed with 10% RPMI-1640. The prepared single cell suspensions were RBC lysed using 1x RBC lysis buffer. 15 × 10^6^ of the bone marrow cells were then re-suspended in 15 mL of 10% RPMI supplemented with 5 ng/mL of GM-CSF and IL-4 and seeded in a 100 mm petri dish for 6 days in a humidified CO_2_ incubator. Half of the culture medium was replaced with complete RPMI supplemented with 5 ng/ml of IL-4 and GM-CSF every alternate day. The cultured cells were harvested after five days using a cell scrapper and washed twice with 10% RPMI at 4 °C for 5 minutes at 200xg. The viability of the in-vitro generated BMDCs was assessed using trypan blue staining. The cells were then transduced to express α-Gal-3 IB as described for the CD8+ T cells in a previous section.

### Western blot and Pull-down assay to confirm the expression of α-Gal-3-IB

The expression of Gal-3 was further confirmed by western blotting and a pull-down assay. The HEK293T cells were transfected with the α-Gal-3-IB and the VC plasmids in the presence of PEI. The transfected cells were collected after 72 hours, scraped off the surface, and stored overnight at -80 °C in C-100 lysis buffer to prepare cell lysates. The next day, the cells in the C-100 lysis buffer were vortexed and centrifuged to obtain a clear supernatant. The clear supernatant or cell lysates were run on SDS-PAGE and transferred to a PVDF membrane, which was then probed with an anti-c-Myc monoclonal antibody. The cell lysates were also used to perform a pull-down experiment to confirm the specificity of the IB for Gal-3. 150 μg/ml Gal-3 protein was immobilized on Ni-NTA beads.

After washing, equal amounts of cell lysates were added to the immobilized beads. The interacting proteins were eluted with 400 mM imidazole, and the eluates were run on SDS-PAGE and transferred to a PVDF membrane. The PVDF membrane was probed with anti-6x His monoclonal antibody and anti-c-Myc monoclonal antibodies. The IB specifically interacted with the immobilized Gal-3, as bands corresponding to Gal-3 and IB (i.e., 30 kDa and 47 kDa) were visible after the membrane was developed with the specific antibodies. To confirm the expression of α-Gal-3-IB in the transduced CD8^+^ T cells, western blotting was performed on the cell lysates from the sorted transduced cells and untransduced cells, and they were probed with anti-c-Myc monoclonal antibody.

### Confocal microscopy

Spleen and lymph nodes were pooled from C57BL/6 mice in 10% RPMI and subjected to RBC lysis after cell suspension was prepared. After RBC lysis, the cells were centrifuged at 1380 rpm for 15 minutes at 4 °C, and the pellet was dissolved in the media. CD8^+^T cells were MACS-purified using the provided protocol. After MACS, the purity of the CD8^+^T cells was assessed. These cells were then spin-transduced in 96-well U-bottom plates using a pseudovirus expressing an anti-Gal-3 sdAb (IB) or the vector control (VC). Spin transduction was carried out for 90 minutes at 800g and 37 °C. The next day, the media was replaced with 10% RPMI, and after 72 hours, GFP-positive CD8^+^T cells were analyzed and sorted for confocal microscopy to assess co-localization of Gal-3 with α-Gal-3 IB or ZAP70 at the immunological synapse.

The coverslips were coated with 1% gelatin overnight, then washed twice with 1xPBS. The next day, they were coated with anti-CD3 and anti-CD28 antibodies and incubated for 30 mins at 37°C. Then, the sorted and transduced cells were added to the anti-CD3 and anti-CD28-coated coverslips and fixed with 4% paraformaldehyde (PFA) for 10 minutes at room temperature. After fixing, the cells were permeabilized with 0.1% Triton for 15 minutes and were blocked with 2% BSA for 1 hour at RT. After washing, the Fc block reagent was added for 1 hour at RT, followed by incubation with the primary antibodies against Galectin-3, ZAP-70 (PE), or Alix in the permeabilization solution for 2 hours. The coverslips were then washed 3 times with 1xPBS. This was followed by 1 hour of incubation with a cocktail of Hoechst and a fluorescent-tagged secondary antibody, anti-Rabbit IgG-Alexa Fluor 647. After incubation, the cells were washed 3 times with 1xPBS and mounted on slides for image acquisition. The images were acquired using the Zeiss Confocal Microscope at 40x or 63x magnification. All captured images were analyzed using ImageJ.

### Viral infection of mice

CD8^+^ T cells were MACS purified from pooled lymph nodes and spleens of OT1xRAG1^-/-^ mice. Control or the α-Gal-3 IB expressing OT1 cells (CD45.2^+^) were transferred into CD45.1^+^ congenic mice. The donor and recipient animals used for adoptive transfer experiments were sex and aged matched. The frequency of the donor cells in circulation was checked before the infection to ensure equal transfer. The animals were then infected with 200pfu of WSN-SIINFEKL via the intranasal (i.n) route. The donor OT1 cells were tracked in circulation prospectively. A heterologous challenge was given i.n with 10^5^ pfu of MHV68-SIINFEKL after three months post primary infection and the recall response was analysed to measure the frequency and number of donor OT1 cells in circulation, lymphoid organs as well as lung tissues at different days post re-infection. For cellular analyses, the mice were bled at different time intervals from the retro-orbital plexus, and the blood was collected in EDTA-containing microcentrifuge tubes. To characterizes cells from lymphoid and non-lymphoid tissues, the mice were sacrificed at different time points. The animals were perfused with 15 mL of 1x PBS. The organs were then collected in 10% RPMI. Lungs were then treated with 1mg/mL collagenase (Type IV) for 30 minutes at room temperature. Collagenase-treated lungs and other organs were processed to form single-cell suspensions. For processing spleen samples, the cell suspensions were treated with RBC lysis buffer and washed twice with 1x PBS.

### Flow cytometric analysis

The blood samples for cellular analysis were processed by incubating the specific antibodies at a 1:500 dilution in 30μL of the blood samples for 30 minutes at 4 °C in the dark. Later, the blood samples were lysed with the RBC lysis buffer (pH 7.3) for 5 minutes at RT. The samples were then centrifuged, and the pellets were resuspended in PBS. The samples were analyzed using a BD FACS Aria Fusion or a BD C6 Accuri.

For staining, the requisite amounts of cell suspensions were collected from different organs across different groups and incubated with antibody cocktails at a 1:500 dilution for 30 minutes. After staining, the samples were washed, and the cells were resuspended in 1x PBS. For intracellular staining, cells were surface-stained, then fixed with fixation buffer. After fixation, the cells were permeabilized with 1x Permeabilization buffer for 1 hour. After permeabilization, the cells were washed and stained with specific antibodies, prepared in the permeabilization buffer for 1 hour. For the intracellular cytokine assay, cells were stimulated with the SIINFEKL peptide for 4 hours at RT in the presence of Brefeldin A. After the incubation, the cells were stained with the respective antibodies. After staining, the cells were washed and resuspended in 1x PBS and acquired using BD FACS Aria Fusion.

### Proliferation and Antigen Presentation Assay

To assess the functionality of the α-Gal-3-IB transduced CD8^+^ T cells, sorted transduced CD8^+^ T cells were labeled with cell trace (Far Red) and they were stimulated with anti-CD3 and anti-CD28 antibodies. The proliferative potential of these cells was compared with that of the Gal-3KO CD8^+^ T cells using Cell Trace dilution, and the frequencies of cells at each division were analyzed. In another set of experiment, BMDCs and CD8^+^ T cells transduced with α-Gal-3-IB and VC were used for antigen presentation assays. Alongside Gal-3KO BMDCs were used for such assays. The sorted control BMDCs, vector transduced and the α-Gal-3 IB expressing BMDCs were pulsed with different concentrations of Ovalbumin or SIINFEKL peptide and were then co-cultured with Cell Trace-labeled CD8^+^ T cells, which were either untransduced or transduced with α-Gal-3-IB and VC. The CD8^+^ T cells from co-cultures were analyzed for their proliferation. ELISA was used for quantifying the levels of Gal-3, IL-2, IFN-γ in the culture supernatants of the cells modified or Gal-3 response by a-Gal-3 IB or those from the co-culture of T cells and BMDCs.

### Quantification of viral titers

To quantify viral titers in lungs infected with MHV68-SIINFEKL, perfused lungs were homogenized in serum-free DMEM and centrifuged at 13000 rpm for 15 minutes at 4 °C to obtain a clear supernatant. Confluent Vero E6 cells were incubated with the lung supernatants at different dilutions. The plaques were observed after 5-7 days and were stained using crystal violet (56,57).

### Statistical analysis

Student’s “t” test, One-way or two-way ANOVA tests were applied as indicated to compare the responses between groups, and the results are presented as Mean ± SD. The p values are shown in the figures or figure legends and are represented as *p ≤ 0.05, **p ≤ 0.01, ***p ≤ 0.001, and ****p ≤0.0001, as indicated in the respective figure legends.

## Funding

This work was supported by extramural grant number IPA/2021/000136 from DST-SERB and ANRF/ARG/2025/000452/LS from Anushandhan National Research Foundation to SS.

## Acknowledgments

We thank the Council of Scientific and Industrial Research (CSIR) for granting the fellowship to Yuviana J. Singh.

## Supporting Information

Fig S1. Cloning of anti-Gal-3-intrabody (α-Gal-3-IB). A-C. Anti-Gal-3 sdAb sequence was cloned into a mammalian expression vector, pLenti-GFP, using the *Xba-1* and *Bam-H1* restriction sites. The sdAb was cloned upstream of the GFP, and a c-Myc tag was inserted at the C-terminal of the antibody sequence during cloning. A. The vector map demonstrates the cloning of the sdAb sequence. B. The amplified sequence of the anti-Gal-3 sdAb using the sdAb-specific forward and reverse primers is shown. C. The amplicon was digested using specific restriction enzymes and then ligated with the pre-digested pLenti-GFP vector. The ligated product was transformed into the Stbl-3 strain of *E. coli*. The positive colonies were screened and confirmed by restriction digestion with *Sal-1* and *Nde-1*. The agarose gel shows the digested product, confirming the cloning in the pLenti-GFP vector.

Fig S2. Assessing the functionality of OT1 cells expressing α-Gal-3-IB. A. The splenocytes were transduced with the control pseudoviruses or the pseudoviruses encoding α-Gal-3-sdAb. After spin transduction and incubation for 16 hours, the splenocytes were activated with different concentrations of the SIINFEKL peptide (0-1μg/ml). After 48 hours of activation, Gal-3 production was assessed. B. Representative FACS plot and offset histograms show the percentage of GFP^+^CD44^+^ cells and Gal-3^+^GFP^+^CD44^+^ cells, respectively. A vertical solid line in the histograms separates Gal-3^+ve^ from Gal-3^-ve^ populations. C. The corresponding bar graphs show the percentage of Gal-3-producing cells in the vector control and IB group. (D) The levels of secreted Gal-3 in the culture supernatants of the stimulated cells were assessed by ELISA, and the absorbance values for the detection of extracellular Gal-3 are shown. The level of statistical significance was calculated using One-way ANOVA. ****p<0.0001, ***p<0.001, **p<0.01, *p≤0.05 and ^ns^p >0.05.

Fig S3. Analyzing the activation and proliferative response of Gal-3KO and α-Gal-3 IB expressing OT1 cells. A-B. CD8^+^ T cells from WT and Gal-3KO mice or the control OT1 transductants and α-Gal-3 IB expressing OT1 cells were stimulated with 1μg/ml of anti-CD3 and anti-CD28 antibodies. The expression of CD44 and CD69 was analyzed at different time points. Bar graphs show MFI values for CD69 and CD44 expression. The fold change values in the Gal-3KO and α-Gal-3 IB groups were calculated by dividing the values for Gal-3KO and α-Gal-3 IB cells by those of the control groups (WT and vector control, respectively). C and D. Representative offset histograms, along with their unstimulated controls, at different time points show the proliferation of WT and Gal-3 KO CD8+ T cells, as well as vector control and α-Gal-3 IB-expressing cells. The dotted vertical lines were used to mark dividing cells. The cell frequencies in each division are shown in bar graphs (C). The values were plotted as Mean ± SD, and the experiments were performed 3-4 times. Two-way ANOVA was used to calculate statistical differences. ****p<0.0001, ***p<0.001, **p<0.01, *p≤0.05 and ^ns^p>0.05.

Fig S4. Comparing the phenotypic profile of Gal-3 KO BMDCs and α-Gal-3 IB transduced BMDCs of WT mice. A-F. The BMDCs were generated from the bone marrow cells of Gal-3KO and WT mice by culturing in RPMI supplemented with GM-CSF and IL-4 for 6 days. BMDCs from WT mice were spin-transduced with either control pseudoviruses or those encoding the α-Gal-3 IB sequence. These BMDCs were compared for the expression of MHC-I, MHC-II, and the co-stimulatory molecules, CD80 and CD86. A and D. Representative offset histograms show the frequency of cell populations scoring positive for the indicated marker from WT and Gal-3 KO mice or the vector control and the α-Gal-3 IB expressing cells. B and E. Bar graphs display cumulative data for the cells positive for the indicated markers. C and F. Bar graphs show the MFI values for the indicated markers obtained from the cells of different groups. Mean ± SD values are plotted. The experiments were performed three times. Two-way ANOVA was used to calculate statistical differences. ****p < 0.0001, ***p < 0.001, **p < 0.01, *p ≤ 0.05 and ^ns^p>0.05. G-H. Representative FACS plots show the pre- and post-sort analysis of BMDCs from different groups. The sorted cells were used in antigen-presenting assays.

Fig S5. Schematic for experiments for assessing *in vivo* differentiation of antigen-specific CD8 T^+^ cells expressing α-Gal-3 IB and the characterization of the transferred cells. A. 40,000 control OT1 cells or the α-Gal-3 IB expressing OT1 cells (CD45.2^+^) were adoptively transferred into the sex-matched congenic CD45.1 mice, which were then intranasally infected with 200 pfu of IAV (WSN-SIINFEKL), and the infected animals were divided into different groups. The infected animals were sacrificed at 7- and 45-days post-infection (dpi), for cellular analysis in lymphoid and non-lymphoid organs. To assess the recallability of memory cells, the animals were intranasally infected with MHV68-SIINFEKL at 90 dpi, and the expansion of the transferred vector-transduced and IB-expressing cells was analyzed at 7 days post-rechallenge. B-C. Characterization of donor cells prior to adoptive transfer. The transduced CD8^+^ T cells were also stained with PI to assess viability. Frequencies of PI-positive cells in both groups are shown in the FACs plots (B) and a bar graph (C). The frequencies of the PI-positive cells are plotted as Mean ± SD. The experiments were performed 3 times. Student’s t-test was used to calculate statistical differences, ****p<0.0001, ***p<0.001, **p<0.01, *p≤0.05, and ^ns^p>0.05. D. The FACs plots represent the pre-sort and the post-sort CD8^+^ T cells in both groups. E. The frequencies of transferred donor cells were measured in recipient blood before primary infection to ensure equal transfer of the vector control and IB-expressing OT1 cells into the congenic mice. The FACS plots and bar graphs depict the frequencies of donor cells in circulation in both groups. Two-way ANOVA was used to evaluate statistical differences. ****p < 0.0001, ***p < 0.001, **p < 0.01, *p ≤ 0.05 and ^ns^p>0.05.

Fig S6. Quantification of antigen-specific vector transduced and α-Gal-3 IB expressing CD8^+^ T cells in different organs at acute (7dpi), memory (45dpi), and after recall phase (97 dpi). A. The frequencies of the donor cells were evaluated in lymphoid organs and lungs at 7 and 45 dpi, as well as at 7 days post recall (dpr) with MHV68-SIINFEKL. B. Representative FACs plots depicting the frequencies of virus-specific donor (CD45.2^+^CD44^+^) cells in the recipients of vector control and α-Gal-3 IB expressing OT1 cells during different time points in lymphoid organs. C-D. Absolute counts of these cells were also plotted at different time points. The experiments were repeated two more times with similar results. The values are represented as Mean ± SD. Two-way ANOVA was used to evaluate statistical differences. ****p < 0.0001, ***p < 0.001, **p < 0.01, *p ≤ 0.05 and ^ns^p>0.05.

Fig S7. Quantification of IFNγ^+^, TNFα^+^, and IFNγ^+^ TNFα^+^ DP cells in different organs at different time points. A. The functionality of the virus-specific VC and α-Gal-3 IB transduced CD8^+^ T cells in the spleen, Med LN, and lungs during different phases was measured by ICCS assays. The cells were stained using a standardized protocol, and the numbers of IFNγ^+^, TNFα^+^, and IFNγ^+^TNFα^+^ DP cells in different organs were quantified. B. Representative FACs plots depict the frequencies of IFNγ^+^, TNFα^+^, and IFNγ^+^TNFα^+^ DP cells in different organs. C-D. Absolute counts of these cells were plotted at different time points. The experiments were repeated three times with similar results. The values were represented as Mean ± SD, and Two-way ANOVA was used to evaluate statistical differences. ****p < 0.0001, ***p < 0.001, **p < 0.01, *p ≤ 0.05 and ^ns^p>0.05.

Fig S8. Quantifying the number of α-Gal-3 IB CD8^+^ T cells in different organs. A-D. After recall of memory cells by infecting animals with MHV68-SIINFEKL, intracellular staining was performed to determine the number of c-Myc positive cells (representing the α-Gal-3 IB expressing cells) in the spleen, MedLN, and lungs. The frequency (B), absolute number (C), and MFI values (C) for c-Myc positive cells were plotted as bar graphs. Mean ± SD are shown, and Two-way ANOVA was used to calculate statistical differences, where ****p<0.0001, ***p<0.001, **p<0.01, and *p≤0.05.

## Notes

### Competing Interest Statement

The authors have declared no competing interest.

